# Are theoretical advances in distance-based tree learning practical for phylogenetic inference?

**DOI:** 10.64898/2026.01.12.698017

**Authors:** Anastasiia Kim, Andrey Y. Lokhov, Marc Vuffray, Ethan Romero-Severson, Emma E. Goldberg

## Abstract

The phylogenetic tree of relatedness for species, viral samples, or other biological taxa is essential information for all fields of evolutionary biology. Many different statistical and computational tools for phylogeny inference are used by practicing biologists, but none take advantage of a body of theoretical work on fast-converging algorithms that guarantee correctness with high probability under particular conditions. Here, we assess the utility of one of the most advanced of these algorithms when applied in reasonable biological situations. Our simulation study shows that realistic datasets will often not meet the assumptions of the algorithm, but also that the results are relatively robust to this problem. We additionally provide guidance on how the algorithm can be deployed when the true tree is not known, which is essential for any real-world application. Overall, our intention is to bring a class of methods to the attention of the phylogenetics community, and to make the algorithmic community aware of the needs of practicing biologists.

## Introduction

Inferring a phylogeny—the tree of relatedness of a set of biological samples—is a crucial step in numerous research fields, from systematics to biogeography to epidemiology to vaccine design [1, 2, 3, 4], and far beyond. Likelihood-based methods are the preferred tool of phylogenetics because of their beneficial statistical properties [5], and an explosion of methods now allow researchers to compute the likelihood of a set of observed DNA sequences under a wide variety of models of sequence evolution and other processes [6, 7, 8, 9, 10]. However, although the theoretical consistency guarantees of maximum likelihood and stationary MCMC chains are very appealing, they are computationally very challenging to obtain in practice [11]. This is particularly true for datasets with a very large number of taxa—such as the millions of genomes obtained during the SARS-CoV-2 pandemic—because the number of possible trees is far too large for a thorough heuristic search of tree space [12].

Consequently, other statistical approaches are of interest. Particularly well-developed are methods based on the matrix of distances between each pair of sequences, which allow direct construction of a tree through clustering algorithms rather than requiring a search of tree space [13]. The distance between two genomes is computed as a single number that summarizes similarities and differences at each site. For biologists, a popular distance-based family of methods is neighbor-joining (NJ). NJ is a bottom-up agglomerative clustering method that builds a tree by iteratively clustering taxa based on a criterion that minimizes total branch length [14]. NJ is statistically guaranteed to reconstruct the correct tree topology if the maximum error in the pairwise distance matrix is less than half the length of the shortest true edge [15]. The more sites that are available in the alignment, the less error is expected to separate the computed distance from the true evolutionary distance. Thus, a key question regarding distance-based methods like NJ is how sensitive they are to limited sequence length. In practice, the strict condition on the distance matrix is frequently not met by real datasets, but NJ is often seen to return the correct tree nevertheless [16, 17]. Although NJ is statistically consistent, it can still suffer from long branch attraction and long branch distraction when sequences are short, which can lead to erroneous clustering of divergent taxa or misplacement due to branch-length variance [18]. Despite this, NJ remains one of the fastest and most practical methods for inferring phylogenies, with algorithmic complexity *O*(*n*^3^) where *n* is the number of taxa. It has particularly been shown to perform well in large-scale analyses where evolutionary rates are moderate and sequence lengths are sufficient [19], when sequence lengths are too short for maximum likelihood to perform well [20], and when the evolutionary rate is relatively constant across the tree [21]. It does, however, in some cases require sequence lengths that increase exponentially with the number of taxa [15, 22, 23]. This may be problematic particularly for pandemic-scale data, in which the number of taxa is orders of magnitude larger than the length of the genome.

Exponential sequence-length bounds, as in NJ, were previously considered the norm for exact topology recovery. A seminal breakthrough came from Erdős et al. [24], who introduced the Short Quartets Method (SQM), which runs in polynomial time in the number of taxa and reliably reconstructs the tree from sequences of length scaling polynomially in the number of taxa—a property later defined as fast convergence [25]. SQM discards large, potentially unreliable distances, instead focusing on building small-diameter subtrees and merging them into a complete topology. Building on this foundation, a series of papers throughout the early 2000s refined the understanding of phylogenetic inference under minimal sequence length required for reconstruction. Several works of Mossel et al. showed that one can reconstruct the tree from sequence length scaling logarithmically with the number of taxa if the mutation probabilities remain below a fixed constant [26, 27, 28, 29], confirming Steel’s “Favorite Conjecture” [30], and otherwise polynomial sequence length is needed [31]. The premise of methods claiming reconstruction from logarithmically scaling sequence lengths becomes very appealing in the scenarios such as viral genomes sampled from a pandemic, where the number of sites in the genome is significantly smaller than the number of taxa.

When some tree branches cannot be reconstructed under the guarantees of any of the above methods, or related ones, the algorithm will either not return any tree, or it will return a complete tree that may contain many incorrect branches [24, 32, 33, 34, 35] (observed by Gronau et al. [25]). To remedy this, subsequent work introduced approaches to reconstruct partial trees (a ‘forest’) with high confidence, instead of one full tree potentially with errors [36, 37]. These methods progressively reconstruct the clearly supported edges of the tree from the taxa inward, stopping once they encounter an indistinguishable edge. Early versions of this strategy allowed shallow indistinguishable edges to block the reconstruction of deeper, statistically supported edges [25], but Daskalakis et al. [38] proposed an algorithm which is able to skip over such edges. An additional feature of Daskalakis et al. [38], relative to all previous algorithms, is that it does not impose fixed bounds on the edge lengths that can be reconstructed. Instead, the user can provide values of hyperparameters that offer a trade-off between resolving short edges or deep parts of the tree.

Despite the appeal of quickly and correctly reconstructing only the parts of a tree that are actually resolvable, these fast-converging forest algorithms are not used within the phylogenetics community, and many questions remain about their performance in biologically-relevant situations. Here, we implement the method in Daskalakis et al. [38]—previously presented only mathematically—and conduct a simulation study to answer questions a potential user would have, such as: How much resolution does a reconstructed forest typically contain? How can one choose appropriate hyperparameter values without knowing the branch lengths of the true tree? How robust is the forest correctness when the data and hyperparameters do not meet the algorithm’s assumptions? Overall, when is the forest approach superior to a simpler distance-based approach such as NJ? Our goals are to introduce the realm of forest algorithms to practicing phylogeneticists, and to illustrate the needs of such practitioners in order to guide the next generation of algorithm development.

## Methods

### The Forest algorithm

We compared results from the original NJ algorithm [14], which produces a single complete tree, against the algorithm of Daskalakis et al. [38] (hereafter ‘Forest’), which reconstructs either a single complete tree or a disjoint forest with multiple components. Note that across all components of a forest each of the taxa is represented once as a leaf, and each component can contain taxa that are either nearby or spread out across the true tree (meaning, that do or don’t comprise one side of a split in the true tree). Both methods work from the matrix of pairwise distances of all sampled taxa. NJ has no tuning parameters, but Forest lets the user control hyperparameters (called *m*, *M*, and *τ*) to determine a trade-off in the resolution of short or deep branches.

Forest works by first building a graph connecting all pairs of taxa whose estimated distance is less than a threshold *m*. Each connected component becomes a local cluster. Within each cluster, the local tree is inferred using a greedy routine called MiniContractor, which detects reliably long internal edges while contracting those too short to resolve (shorter than the threshold *τ*). The size of the local neighborhood used to search for detectable internal edges is defined by the ‘ball,’ which includes all leaves within distance ≤ *M* of a given pair. If *M* is large enough that the ball covers the entire cluster, MiniContractor can directly infer full bipartitions without additional steps. Otherwise, the algorithm proceeds to a final extension phase, assigning the rest of the leaves in the cluster to the appropriate side of each split. The final output is a forest of components, each capturing the reliably-inferable structure within its respective subset of taxa. With high probability, the algorithm reconstructs only branches that are strongly supported by the data; otherwise it contracts branches to produce polytomies, or it omits branches by leaving components unconnected to one another.

The theoretical guarantees of correctness in the forest require that a (*τ, M*)-distorted metric satisfies the ‘distortion condition’: for all ‘short’ pairwise distances (either true or estimated) that are less than *M* + *τ*, the difference between true and estimated pairwise distances must be at most *τ* [37]. The algorithm then reconstructs a forest with chord depth up to ∼ *M/*2 (shallow enough edges) which includes all edges of length at least 4*τ* (edges that are long enough). Therefore, the algorithm provides a trade-off between the resolution of short branches (controlled by *τ*) and the depth of the reconstructed forest (controlled by *M*), with *m* controlling the number of components [38, 25].

The Forest algorithm runs in polynomial time (*O*(*n*^5^) for *n* taxa) for a given set of hyperparameters. Without prior knowledge of the true tree, however, it is impossible to know which combinations of hyperparameter values satisfy the distortion condition. It is also not clear a priori which hyperparameter combinations will produce the best-resolved forest for a given dataset. We therefore performed a search across hyperparameter values on each simulated dataset, to try to balance the resolution of the reconstructed tree with the sequence length requirements.

The Forest algorithm was presented only mathematically in Daskalakis et al. [38]. We implemented it in Python, available at https://github.com/lanl/distphylo. User inputs are a dataset in the form of a pairwise distance matrix, and values for *M*, *m*, and *τ*. If multiple values are given for one or more hyperparameters, output is produced for each parameter combination, allowing systematic exploration of the parameter space to assess how different settings affect the output. The Forest algorithm produces a set of bipartitions, from which we construct trees using a tree-popping algorithm [39]. If the bipartitions are conflicting and lead to multiple incompatible trees, no resulting forest is returned.

### Tree inference correctness

For an inferred tree or forest returned by either NJ or Forest, we wish to quantify its correctness relative to the true tree that was used to generate the simulated alignment. We are particularly interested in scoring Forest on its resistance to reporting incorrect information, as well as its power to report correct information.

We use ‘splits’ to score correctness: each internal branch on an unrooted tree defines a split, which is a bipartition of the leaves representing a resolved evolutionary relationship. (We disregard trivial splits, for which an external branch splits one leaf from all others.) The popular Robinson-Foulds (RF) tree distance metric [40] counts the number of splits that differ between two trees. RF is not appropriate for our purposes, however, because it does not define a distance between a single tree and a forest with multiple components. Furthermore, we do not necessarily wish to score an inferred polytomy (a node with degree greater than three) as an error, because we consider it potentially advantageous that Forest can report uncertainty. Therefore, for each forest, we score the number of splits that it defines—where forests with more components and/or more polytomies contain less information and fewer splits—and the proportion of those splits that do not conflict with the true tree. An example is shown in Fig. 1.

**Figure 1:**
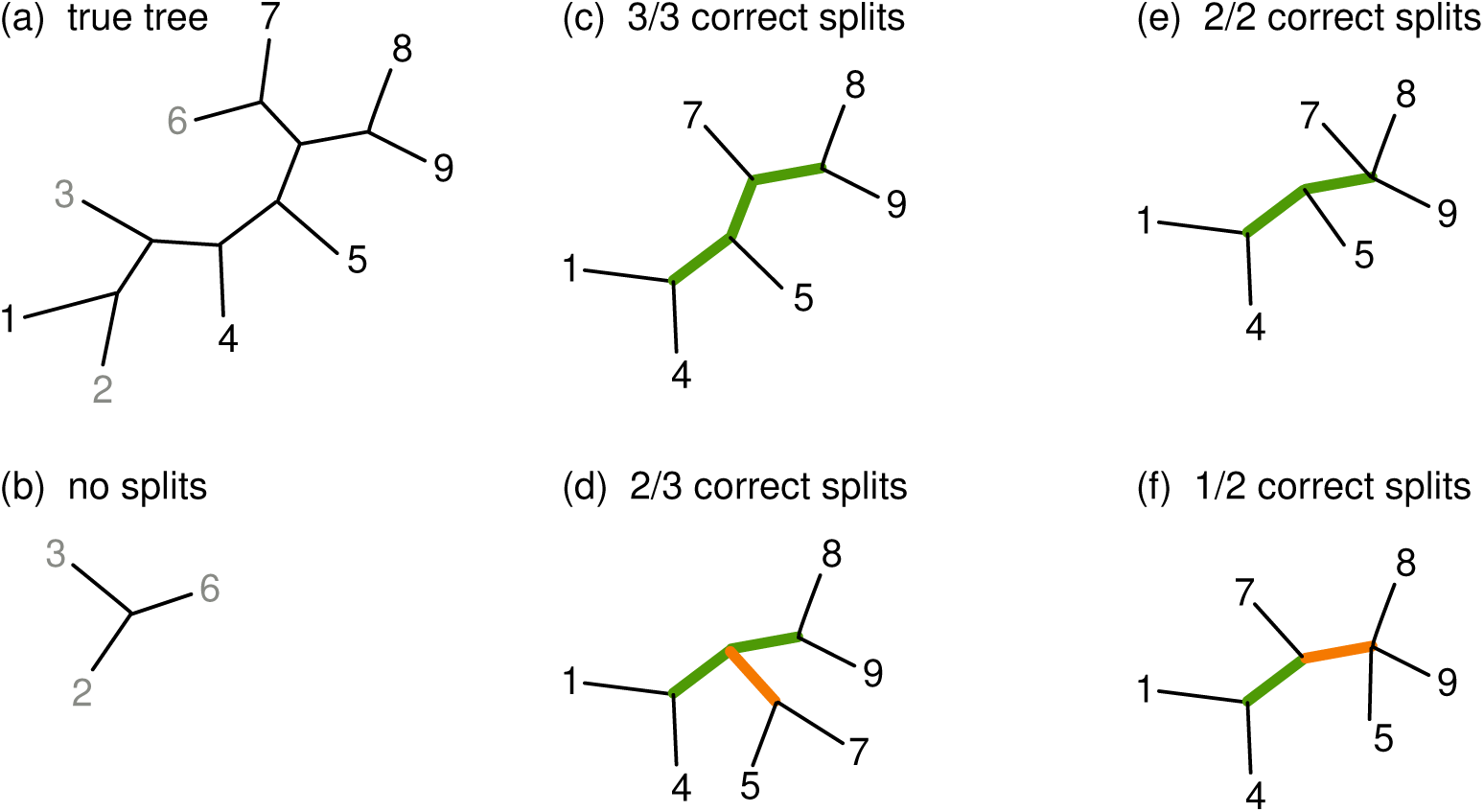
The correctness of forest components is measured by splits defined by internal edges. (a) A true tree is fully resolved, here with *n* = 9 leaves and *n* − 3 = 6 internal edges (when the tree is unrooted) that each define a non-trivial split. Suppose that the inferred forest contains two components, distinguished by gray and black leaf labels. (b) The component with gray labels inherently provides no information because it contains only three leaves and no internal edges. Four possibilities for the component with black labels are shown in panels (c–f). (c) The component is fully resolved, and the three splits it defines are correct, because they are each present in the true tree when it is pruned to these leaves. (d) The component is fully resolved, but it incorrectly defines the split 5, 7|1, 4, 8, 9, which is not present in the true tree. (e) The component has a polytomy and thus only defines two splits, but they are both consistent with the true tree. (f) The component has a polytomy, and it incorrectly defines the split 5, 8, 9|1, 4, 7, which is not present in the true tree.

Based on these scores from splits, we define three metrics for successful tree or forest reconstruction. The first, most stringent metric tests for both reliability and resolution: it requires that a single, fully-resolved tree is produced, and that all of the splits on it are correct. The second metric relaxes the resolution requirement, requiring only that at least 90% of the true splits are found, with the others absent due to multiple forest components or polytomies. It retains the reliability requirement, though, allowing no splits that conflict with the true tree. The third metric tests for reliability only: regardless of the number of splits present, all of them must be consistent with the true tree. This scoring approach allows us to assess how well Forest avoids mistakes (by reporting only true relationships and uncertainty) and how much power it retains.

### Simulated data

To evaluate performance of Forest and compare it with NJ, we conducted a simulation study. Our goal was to identify the minimum sequence length required to produce at least 90 out of 100 successful simulation replicates. Success was scored separately for each of the three metrics (defined above, comprised of reliability and resolution), and separately for trees with different numbers of taxa.

We simulated trees with 10, 20, 30, 50, 70, or 100 leaves, *n*. For each *n*, 20 tree topologies were generated by a birth-death process. On each topology, branch lengths were drawn randomly from a uniform distribution: values between 0.005 and 0.05 define the short branch scenario, and values between 0.05 and 0.1 define the long branch scenario. For each tree, we then simulated replicate sequence alignments under the Jukes-Cantor model using AliSim [41] with rate parameter 1. For each alignment, the pairwise distance matrix was computed also under the Jukes-Cantor model. The length of the sequence alignment, *k*, ranged from 100 to 100,000; it was adjusted by a bisection approach to identify the minimum sequence length needed for success.

Our purpose with this study is to implement and test Forest algorithm under conditions that are somewhat more biologically plausible than what has been done before. We do not use real data because the true tree would not be known, or at least not for sufficiently many datasets. The tree sizes, *n*, we consider are relatively modest, but they are sufficient to show clear patterns in the results, and they present substantial computational burden for Forest. The sequence lengths, *k*, are also only as large as necessary but do cover the range for many viruses or genes. Previous mathematical work has mostly studied the binary Neyman model [34, 26, 42, 29], but we use the simplest sequence evolution model that allows for four nucleotides. We avoid the complication of a mismatch between the generating and inference models of sequence evolution. Importantly, with our mutation rate and branch lengths, the probability that a base (at one site in the alignment) differs from one end of a branch to the other is 0.0037, 0.037, and 0.07 for branch lengths 0.005, 0.05, and 0.1, respectively. These parameter values thus yield alignments with sufficient information to reconstruct a fully correct tree, as evidenced by the NJ results below, and look ‘reasonable’ to the eyes of the authors who work on viral phylogenetics. In comparison, Erdős et al. [24] requires mutation probabilities in the range of 0.1–0.5 for trees of size *n* = 10–1000 to need only polylog sequence length, and Daskalakis et al. [36] conducts simulations for mutation probabilities of 0.1–0.3. These represent very fast mutation rates (or very long tree branches) by most biological standards.

The final, essential piece of biological realism that we include is avoiding the use of any knowledge of the true tree when attempting to infer it from data. This realism has no effect on NJ, other than blinding us to whether its conditions for guaranteed correct inference are met. It does, however, introduce the substantial complication of a grid search across the hyperparameters of Forest. Specifically, for the least-stringent correctness criterion: for the short-branch regime we used *M* ∈ {2.5, 1, 0.5, 0.25}, *m* ∈ {0.6, 0.4, 0.3, 0.2, 0.1, 0.07}, and *τ* ∈ {0.03, 0.01, 0.005, 0.003, 0.0025, 0.00125}; for the long-branch regime we used *M* ∈ {5, 2, 1, 0.5}, *m* ∈ {0.8, 0.6, 0.5, 0.4, 0.3, 0.2}, and *τ* ∈ {0.08, 0.06, 0.05, 0.03, 0.025, 0.0125}. For the other two criteria, to reduce computational time we used a reduced grid with *M* = {5}, *m* = {2}, and *τ* = {0.025} in the long-branch regime, and the same *M* and *m* but *τ* = {0.0025} in the short-branch regime.

In these reduced settings, *m* was chosen sufficiently large to yield a single component, *M* large enough to cover all tips in the tree, and *τ* set to one half of the expected minimum branch length because branch lengths were generated uniformly. All of the above settings were applied to trees with *n* ∈ {10, 30, 50, 70, 100}; for each *n* we sampled 20 random trees and ran at least 100 replicates per tree. Finally, to study parameter effects more closely for *n* = 128, we used a dense grid with *M* ∈ {2.5, 2.4*,…,* 0.5} (step 0.1), *m* ∈ {0.8, 0.7, 0.65, 0.6, 0.55, 0.5, 0.45, 0.4}, and *τ* ∈ {0.01, 0.015, 0.02*,…,* 0.10} (19 values from 0.01 to 0.10 inclusive).

## Results

### Quality of recovered trees and forests

To produce a fully resolved and correct tree (in 90% of the simulation replicates), we found that Forest requires a much longer sequence (larger *k*) than NJ does, especially for more taxa (larger *n*) (Fig. 2). The algorithm can, however, produce most of a correct tree with substantially less sequence data. When requiring only that 90% of the true splits are found, with no incorrect splits reported, Forest requires less sequence length than is needed for the fully-correct tree (Fig. 3). This is particularly true for some replicates, as evidenced by the upper data points in Fig. 2 compared with those in Fig. 3. In essence, final branch resolution is often the most data-demanding step for Forest. Even when only 90% resolution is required, though, Forest still requires greater sequence length than NJ does, especially for larger numbers of taxa (Fig. 3).

**Figure 2:**
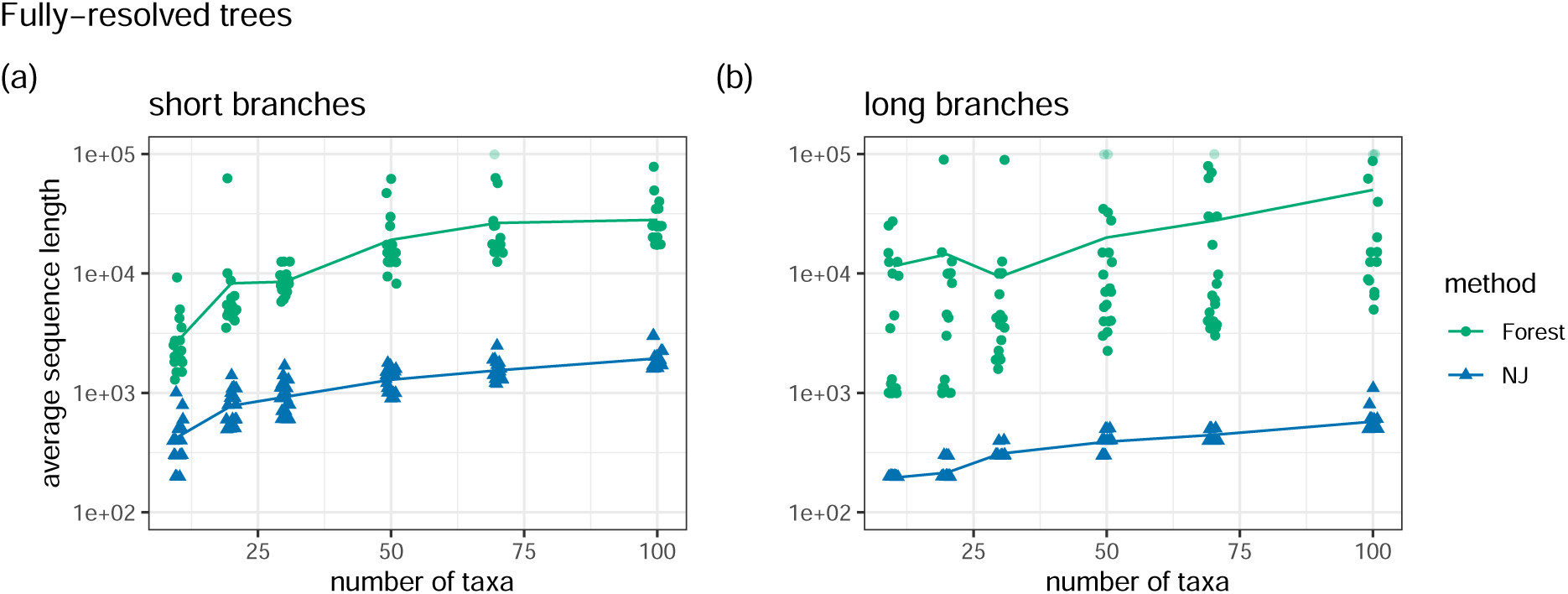
Reconstruction of a fully-resolved tree, with no incorrect splits. Each point shows results averaged across 100 sequence simulation replicates on one true tree. The same tree topologies are used in the two panels, with assigned branch lengths from either the (a) ‘short’ or (b) ‘long’ distribution. Tree size, *n* is on the horizontal axis, jittered to reveal overlapping points. Solid lines connect the average values for each tree size. The average minimum sequence length, *k*, needed to obtain a correct result on 90% of the replicates is on the vertical axis; note the log scale. Lighter-shaded points (at *k* = 100, 000) indicate that the maximum sequence length tested was insufficient.

**Figure 3:**
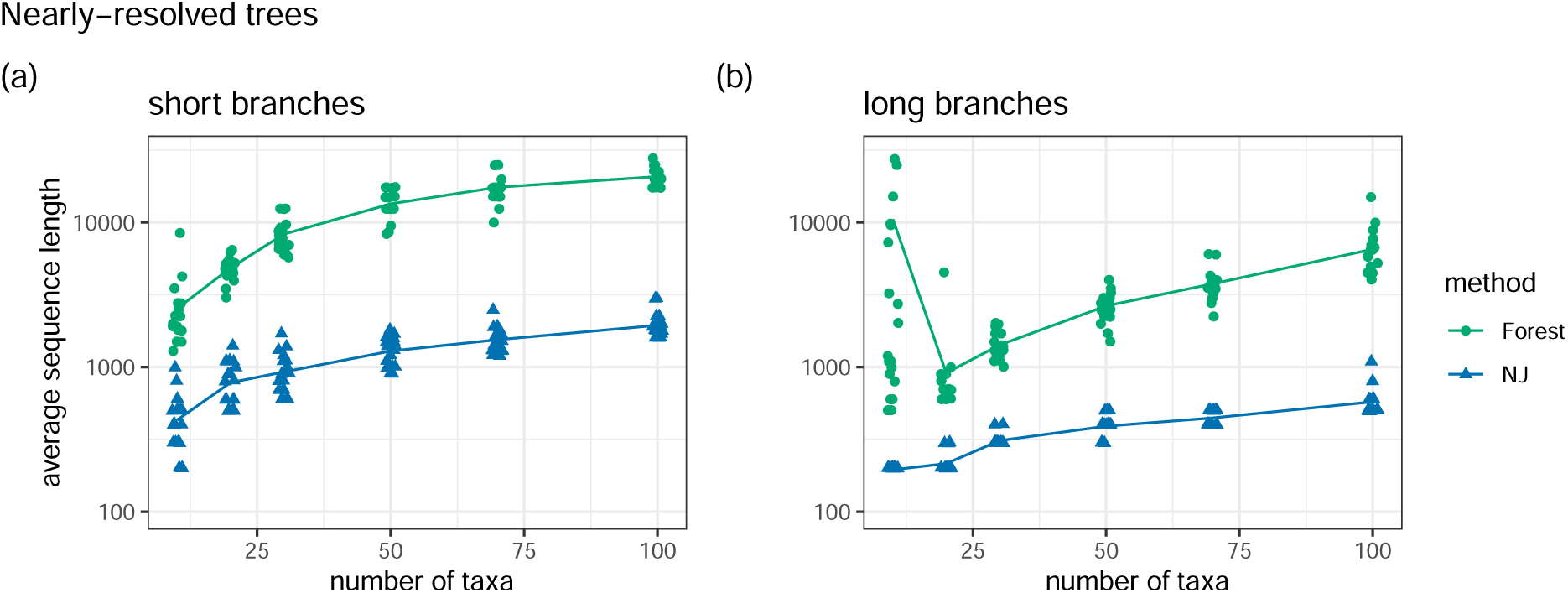
Reconstruction of a nearly-resolved tree or forest, with 90% of the correct splits and no incorrect splits. Figure components are as in Fig. 2, but note the smaller vertical axis scale.

When the requirement is only reliability, not resolution, Forest requires less sequence length to ensure that it returns no splits that contradict the true tree (Fig. 4). On trees with short branches, Forest performs much better than NJ by this metric (Fig. 4a). In the case of long branches, Forest results are similar to those for short branches but NJ results are better, so Forest and NJ perform similarly.

**Figure 4:**
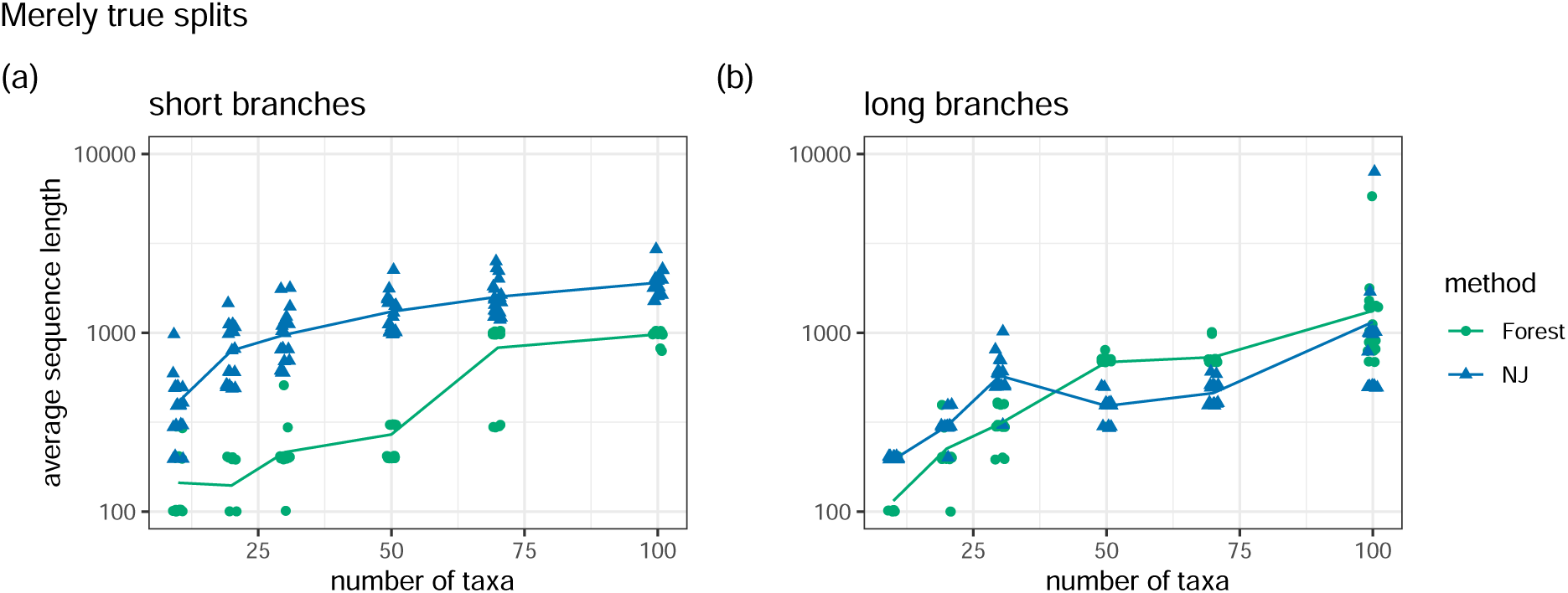
Reconstruction of a partly-resolved tree or forest, with no incorrect splits. Figure components are as in Figs. 2 and 3.

Our three metrics of tree correctness are designed to display the different amounts of resolution that might be present in a forest with polytomies, while retaining reliability. Because NJ always produces a single bifurcating tree, however, the correctness of its output will be the same under all three metrics. Differences in NJ results across Figs. 2–4 are therefore due only to the influence of the Forest results—we keep only replicates for which the Forest algorithm ran to completion, and we prune each NJ tree to the same taxa subsets as in the forest components.

### Effects of hyperparameters

For a biologist with a single dataset, *k* and *n* will be fixed. While the results above may provide an indication of the quality of tree or forest inference that is possible from the alignment, a more practical question is how to choose hyperparameter values that will yield the phylogeny with the most information possible. It is therefore useful to understand how the hyperparameters work together to yield a forest. We illustrate this discussion with a single tree of moderately large size (*n* = 128 taxa) and one realization of either short or long sequence length (*k* = 512 or 100, 000). Results discussed below are shown in Fig. 5. Note also that NJ on the alignment with either *k* yields all 125 correct splits. The general statements made below also hold across several other example datasets that we examined similarly (results not shown).

**Figure 5:**
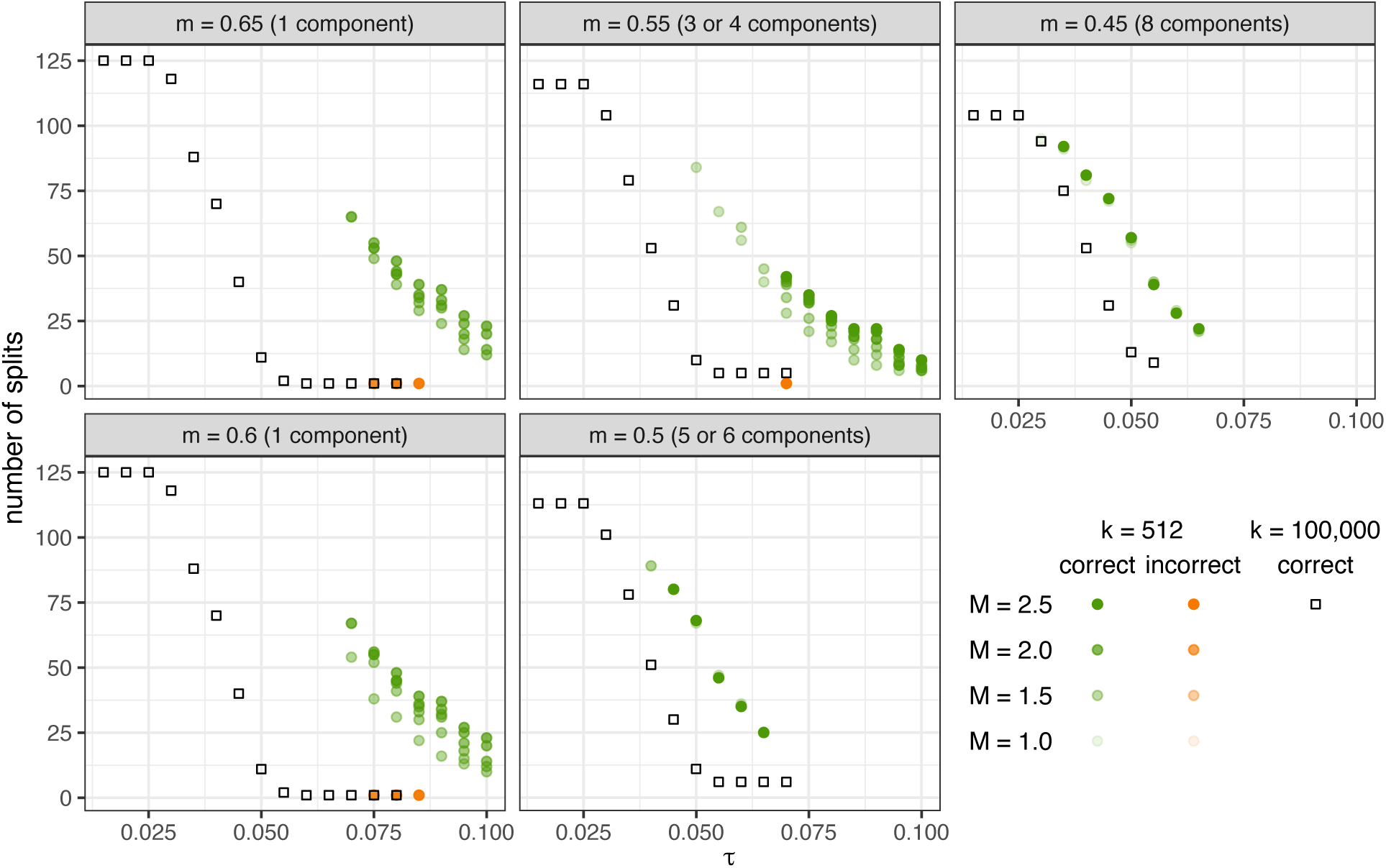
Effects of hyperparameters *τ*, *m*, and *M* on forest resolution and correctness. Results are shown for two alignments simulated on the same tree, with number of taxa *n* = 128. Numbers of splits in the inferred forest are shown for each combination of hyperparameters. For short sequence length, *k* = 512, splits are colored green if correct and orange if incorrect. For long sequence length, *k* = 100, 000, all splits are correct. Values of *M* range from 1.0 to 2.5 in increments of 0.1 and are shown by transparency for short *k*; for long *k* results are nearly identical for all *M* so only *M* = 2.5 is plotted.

The choice of *τ* is restricted by the distortion condition of the Forest algorithm, which states that for all pairwise distances observed from the data or taken from the true tree that are less than *M* + *τ*, the maximum absolute difference between the observed and the corresponding true distances should be less than *τ*. Satisfying this condition is tricky in practice, when the user does not know the true tree and likely does not have a reliable estimate of the minimum branch length. For very large *k*, yielding very small distance errors, the user can safely push *τ* to small values for best results. We saw this with large *k* dataset, in which the distortion condition was met for all but the smallest values of *τ* we considered (the maximum proportion of pairwise distances that violate the distortion condition was 5% for *τ* = 0.015). For smaller *k*, however, it is important to know how robust the Forest results are to violation of the distortion condition. In our example with small *k*, the distortion condition was always violated, by 30–73% of the pairwise distances for the largest and smallest values of *τ*, respectively, that we considered. Consequently, returned forests did contain incorrect splits. However, we found that the number of incorrect splits was low, and it did not worsen as *τ* was decreased to obtain more correct splits. For even smaller values of *τ*, a different kind of output was obtained: the algorithm returned a set of bipartitions that were not self-consistent and so could not be assembled into a unique forest. Therefore, it appears that Forest results are rather robust to values of *τ* that are small enough to violate the distortion condition—few incorrect splits will be obtained or the algorithm will simply not run.

Hyperparameter *M* can be chosen to be sufficiently large to meet the distortion condition. We found that, for given values of *τ* and *m*, larger values of *M* generally yielded more correct splits for small *k*, and had essentially no effect for large *k*. The algorithm reconstructs forests with chord depth on the order of *M/*2, so larger values of *M* generally allow deeper portions of the forest to be inferred. Overall the effect of *M* seems to be minor, and choosing a good value does not seem to be a concern.

Hyperparameter *m* is not involved in the distortion condition, though it must be chosen to satisfy *m <* 1*/*2(*M* − 3*τ*) and *m >* 3*τ* in order for the algorithm to run. When *m* is larger than the chord depth of the true tree, the forest has a single component. Smaller values of *m* yield forests with more components. Having fewer components in a forest might seem advantageous, because more splits can then in principle be obtained (up to the limit of a single fully-resolved tree). For large *k*, we indeed saw that more splits were obtained when there were fewer components (for the best forests with small *τ*), up to all 125 correct splits with one component. However, for small *k*, we found that smaller values of *m*, which create more components, can sometimes allow smaller values of *τ* to produce a valid forest, and thus yield more total splits. In practice, then, for datasets with small *k*, exploring values of *m* is worthwhile.

## Discussion

Based on the proven scalings, the Forest algorithm of [38] in principle allows parts of a phylogeny to be correctly reconstructed with shorter sequence data, compared with other methods that require longer sequences or may include errors in the reconstructed tree. To assess the possible utility of this algorithm for practicing biologists, we tested its performance in more realistic scenarios than it has previously been challenged with.

As expected, we found that Forest outperforms NJ in terms of requiring shorter sequence length to avoid returning a tree/forest that contains any errors when branch lengths are relatively short (Fig. 4a). We also found that Forest performs worse than NJ in many other situations. It does no better at avoiding errors when branch lengths are longer but still reasonable by biological standards (Fig. 4b), and it is much slower than NJ. To produce a correct fully- or mostly-resolved tree/forest, Forest requires much longer sequences than NJ does (Figs. 2 and 3). Furthermore, without prior knowledge of the true tree, it is impossible for a user to know whether the hyperparameter settings for Forest meet the requirements of the algorithm, and thus whether the returned forest is indeed free of errors. This defeats one of the most appealing aspects of Forest: that it returns only the portions of the true tree that can be correctly inferred.

However, if a practitioner is willing to accept that the inferred forest may contain a small number of errors, we found that Forest results are fairly robust to violation of its assumptions (Fig. 5). When hyperparameter *τ* is too small to meet the distortion condition, the returned forest still has no or only few incorrect splits, or else the algorithm simply doesn’t yield an answer. We also found that a user can identify good hyperparameter values without a computationally-expensive grid search. The branch lengths from the NJ tree could be combined with the known value of *k* to approximate the likely error between the true and observed pairwise distances, which would inform a range for *τ*. The chord depth of the NJ tree could also indicate a reasonable starting value for *m*. Overall, if a user is willing to tolerate a small number of incorrect splits, we found that the best Forest results are obtained by finding the minimum value of *m* that yields one component and then trying slightly smaller values, setting *M* to more than 2*m* (it must be at least 2*m* + 3*τ*), and in each case finding the smallest value of *τ* for which the algorithm will run. Note, however, that the resulting forest is still often less resolved and less correct than the NJ tree.

### The potential utility of forests in phylogenetics

Within the realm that Forests performs better than NJ, or if forest algorithms can be improved in the future, biologists could use forest results in several ways. If the forest omits uncertain deep branches (which tend to be long) but includes confident shallow branches (which tend to be short), it effectively reconstructs clades. This could be particularly useful for epidemiology or taxonomy, for example. Pathogen samples obtained during an epidemic may tend to be closely related, such as when limited samples are obtained through contact tracing, or when modern cheap and rapid sequencing of viral genomes enables entire epidemics to be densely sampled. Forest methods could perform relatively well on such data. Or, for public health studies with the goal of identifying recent transmission clusters, forest methods that return clades would be just as useful as phylogenetic methods that return entire trees. For taxonomy, if forest methods tend to recover shallow clades in the tree, they would be particularly useful for refining relationships within a species or genus. Alternatively, if forest components tend to contain tips that do not each form a monophyletic group, they could be more useful for obtaining deeper relationships, which would be most applicable when taxon sampling is strategically over-dispersed. Thus, systematically assessing whether forest components tend to be clades would be useful to understand the areas of biology that would find such methods most useful. We did not see clear such tendencies, but better tests would need to be run on trees with more realistic branch length distributions (e.g., generated by a birth-death process rather than sampled from a uniform distribution).

For forest methods to become popular and useful in biological applications, existing methods for downstream inference (learning about biology from reconstructed phylogenies) would need to be modified to work on forest components. This could include methods that fit single models across multiple components, or that analyze each component and combine the results. Such component-based approaches would be expected to be more reliable than whole-tree approaches when forests contain fewer incorrect splits. The trade-off would be if forests are substantially less resolved than the correct portions of complete trees, which we did often see in our results. A possible hybrid solution for future investigation would be whether forest methods can be used to identify subsets of tips for which reliable reconstruction is expected, but then better resolution within each component can be obtained by NJ or other whole-tree methods.

Alternatively, if forest components are reliable but whole trees are still needed for downstream analyses, another hybrid approach would use forests as inputs into algorithms that merge portions of a tree into a single unified topology. For example, Disjoint Tree Merger methods can either blend components that are not clades or more simply connect components that are [43, 44, 45]. Or a third hybrid option could be to use forest components as topological constraints during the search for a maximum likelihood tree, or as constraints or priors during Bayesian inference; this is possible with standard phylogenetics software [7, 8, 9, 10].

In conclusion, we see great benefit to the phylogenetics community of methods that return a reliable forest, which omits portions of the tree that cannot be reconstructed correctly. Such methods could help practitioners understand which parts of their data are informative, as well as yield downstream results with potentially greater accuracy and more appropriate uncertainties. The currently-best Forest method only delivers results superior to NJ in very limited circumstances, but perhaps future development will lead to improved algorithms. Additionally, it remains untested whether Forest is more robust than NJ in the situation where sequences truly evolve under a different model than is used to compute the pairwise distance matrix. Robustness to this type of model mismatch would be of great value to biological studies, in which the reality of sequence evolution can be extremely complex, and distance-based methods have been shown to be more error-prone than maximum likelihood [46].

## Acknowledgements

Research presented in this article was supported by the Laboratory Directed Research and Development program of Los Alamos National Laboratory under project number 20240198ER. AYL acknowledges support from the U.S. Department of Energy/Office of Science Advanced Scientific Computing Research Program. This research used resources provided by the Los Alamos National Laboratory Institutional Computing Program, which is supported by the U.S. Department of Energy National Nuclear Security Administration under Contract No. 89233218CNA000001.

